# Fundamental Sex Differences in Cocaine-Induced Plasticity of D1- and D2-MSNs in the Mouse Nucleus Accumbens Shell

**DOI:** 10.1101/2023.06.28.546959

**Authors:** Andrew D. Chapp, Chinonso A. Nwakama, Pramit P. Jagtap, Chau-Mi H. Phan, Mark J. Thomas, Paul G. Mermelstein

**Author notes:** Equal contribution (Andrew D. Chapp, Chinonso A. Nwakama). Correspondence to: Mark J. Thomas, Paul G. Mermelstein.

## Abstract

Cocaine-induced plasticity in the nucleus accumbens shell of males occurs primarily in D1 dopamine receptor expressing neurons (D1-MSNs), with little if any impact on D2 dopamine receptor neurons (D2-MSNs). Using *ex vivo* whole cell recordings in male and female mice, we observe alterations in D1-MSN excitability across the estrous cycle similar in magnitude to the actions of cocaine in males. Furthermore, cocaine shifts estrous cycle-dependent plasticity from intrinsic excitability changes in D1-MSNs to D2-MSNs. Overall, while there are similar cocaine-induced disparities regarding the relative excitability of D1-MSN versus D2-MSN between the sexes, in males this is mediated through reduced D1-MSN excitability, whereas in females it is due to heightened D2-MSN excitability.

## MAIN

Cocaine alters the neurophysiology of medium spiny neurons (MSNs) of the nucleus accumbens^1–7^, with enduring drug-induced plasticity observed in the shell (NAcSh) subregion^2^. In male rodents, cocaine reduces the firing rates of D1 dopamine receptor expressing neurons (D1-MSNs), with no effect on D2 dopamine receptor expressing neurons (D2-MSNs)^8–10^. Regarding cocaine-mediated glutamatergic synaptic plasticity, various groups have reported some degree of change at D1-MSN characterized by alterations to miniature excitatory postsynaptic current (mEPSC) frequencies, and/or amplitudes^8, 11, 12^. Similar studies using females have been sparse and data have often been pooled with male subjects^11, 13-15^, most likely because it has been widely assumed these fundamental effects of cocaine would not differ between the sexes. Indeed, in subtype-unidentified nucleus accumbens MSNs, glutamatergic synaptic plasticity following cocaine was reported to be sex-independent^16^. Yet, by isolating D1-MSNs from D2-MSNs and sufficiently powering comparisons between sexes, we report both estrous cycle effects in MSN excitability and major sex differences in response to cocaine, with drug exposure producing a switch from D1-MSN to D2-MSN plasticity, dependent upon the stage of the estrous cycle.

Various genetic backgrounds can produce large sex differences in cocaine-mediated locomotor sensitization^17^. Here we used a mouse line where the actions of cocaine on male and female locomotion were extremely similar, thus potential differences in NAcSh D1-MSN and D2-MSN physiology could not be attributed to differences in behavior. Studies followed previous methods, using a standard psychomotor sensitization protocol followed by 10-14 days of abstinence and then *ex vivo* whole cell current- or voltage-clamp recordings of the NAcSh (Fig 1a)^2, 17^. The estrous cycle was also monitored prior to electrophysiology recordings in order to compare responses during diestrous (before hormonal surges) and estrous (after)— periods within the estrous cycle where the largest differences in MSN physiology^18^ and behavior^19^ have been reported.

**Fig 1.**
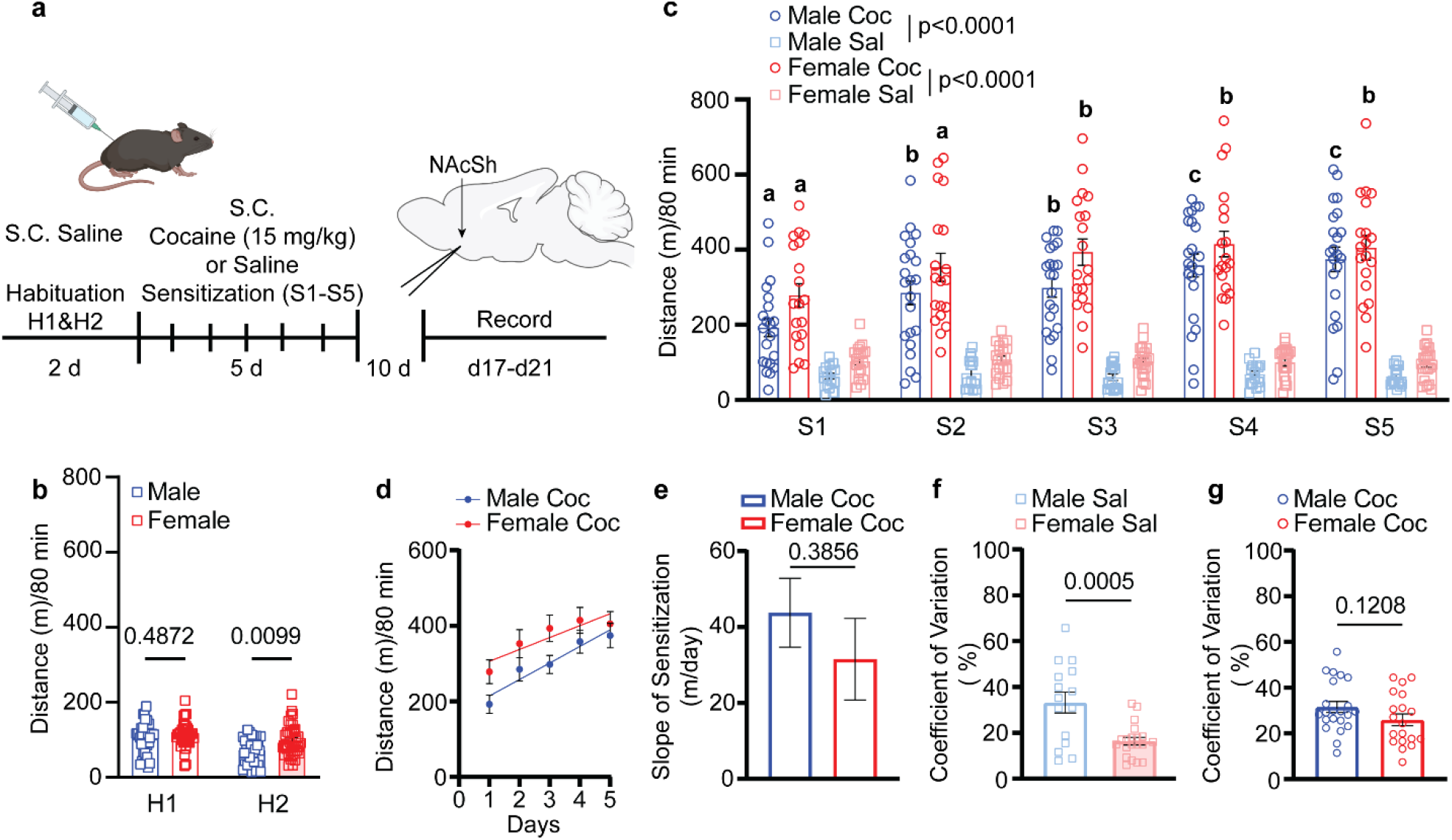
Cocaine psychomotor sensitization in male and female mice. **a**, Experimental timeline. **b**, Two-day saline habituation (H1 & H2) summary data of grouped male (n = 37) and female (n =40). **c**, Psychomotor sensitization, letter symbol “b” significantly different from “a” within sex, “c” significantly different from “a” and “b” within sex, (cocaine: male (n=22), female (n=20); saline: male (n=15), female (n=20). **d**, Linear regression of cocaine psychomotor sensitization. **e**, Slope of locomotor sensitization. **f**, Coefficient of variation for saline-treated animals. **g**, Coefficient of variation for cocaine-treated animals. Abbreviations: Saline (Sal), cocaine (Coc), subcutaneous (S.C.).

Following two days of habituation where all mice (n=77) received saline in the activity chamber (Fig 1b), mice were subdivided, receiving either daily cocaine (15 mg/kg) or saline for the next five days (Fig 1c). Cocaine increased the total distance traveled in both male (n=22) and female (n=20) mice compared to their saline (n=15 male, n=20 female) counterparts. Sensitization to cocaine was observed in males starting on day two of cocaine treatment; day three for females. By day five of cocaine administration, both males and females were exhibiting similar locomotor responses. Of note, on day two of habituation, females displayed a small but significant increase in total distance traveled relative to male counterparts when given saline, a trend that continued throughout subsequent days of testing in the saline-treated controls. We also examined whether there were any sex differences in cocaine sensitization by analyzing the slope of total distance traveled on days one through five (Fig 1d). This analysis represented the average increase in distance traveled per day within sex relative to the previous day^17^. We found no difference in the magnitude of sensitization between males and females (Fig 1e), again indicating minimal sex differences in the psychomotor response to cocaine in this line of transgenic mice.

For the last component of the behavioral analysis, we measured the coefficient of variation for each animal across the 5-day behavioral testing with respect to distance traveled and compared the coefficient across sexes. In saline-treated animals, females had a reduced coefficient of variation relative to their male counterparts (Fig 1f). This finding is in line with a previous report showing that exploratory behavior was more consistent among female than male mice, regardless of estrous cycle^20^, and that females do not increase behavioral variability^21^. Interestingly, cocaine exposure normalized the coefficient of variation between sexes (Fig 1g), a finding consistent with what we observed in various other mouse strains^17^. Overall, the data indicate similar locomotor sensitization following cocaine in these male and female mice.

Next, we investigated cocaine-induced changes to excitability and glutamatergic synaptic physiology in the NAcSh among D1-MSNs and D2-MSNs during late abstinence^2^. We recorded from a total of 302 neurons. As others have reported^8–10^, D1-MSN excitability is reduced relative to D2-MSN excitability in saline-treated males (Fig 2a). Cocaine exposure followed by drug abstinence led to a more pronounced reduction in the firing frequency of D1-MSNs in males (Fig 2b). No change was observed in male D2-MSN excitability following cocaine treatment (Fig 2c). The net effect of cocaine in males is an augmented gap between D1-MSN and D2-MSN excitability, driven by reduced D1-MSN activity (Fig 2d,e).

**Fig 2.**
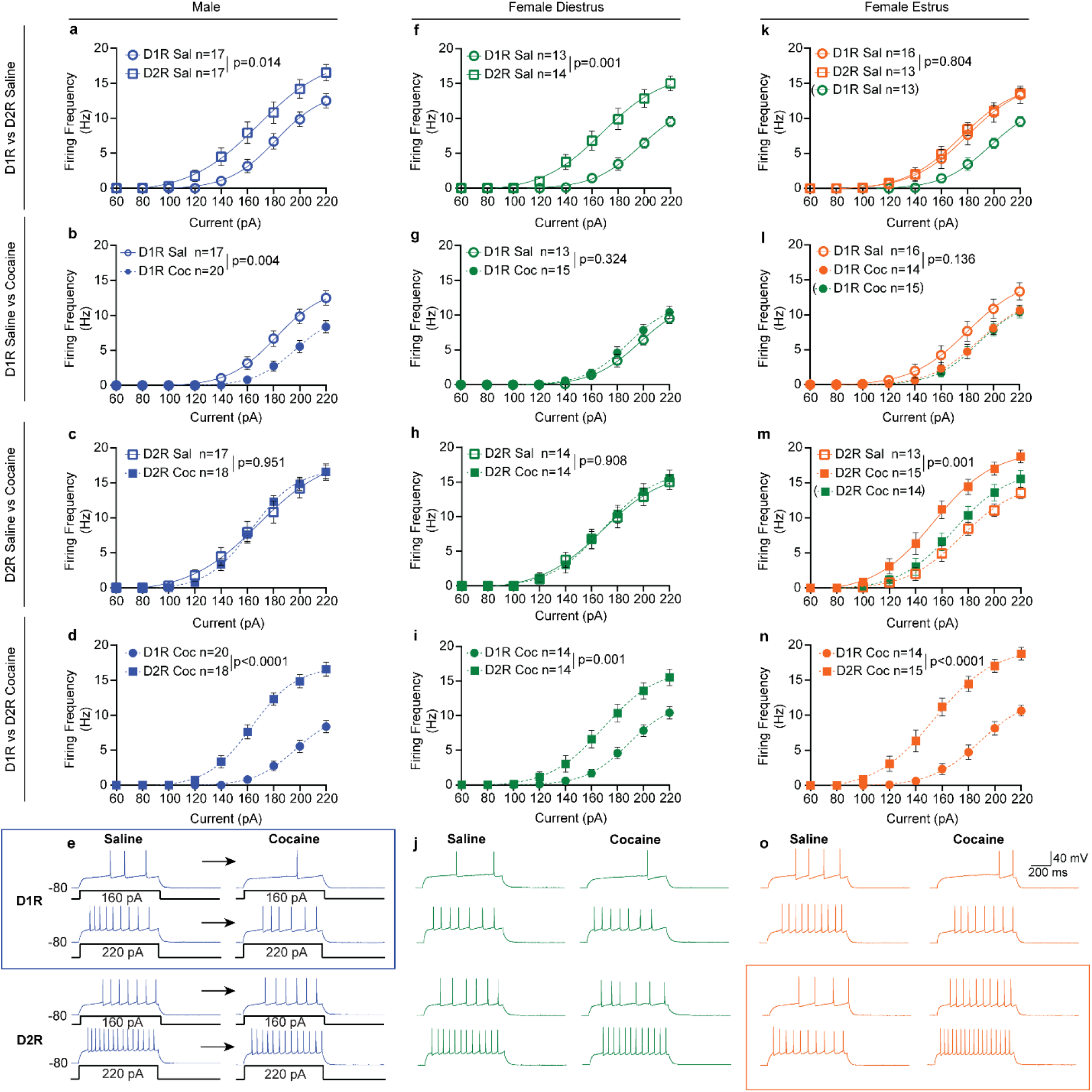
NAcSh neuronal excitability in male and female mice. **a**, Summary data for saline-treated male D1-MSN vs D2-MSN neuronal excitability. **b**, Summary data for saline-treated vs cocaine-treated male D1-MSN neuronal excitability. **c**, Summary data for saline-treated vs cocaine-treated male D2-MSN neuronal excitability. **d**, Summary data for cocaine-treated male D1-MSN vs D2-MSN neuronal excitability. **e**, Representative raw traces from saline-treated and cocaine-treated male D1-MSNs (top), and D2-MSNs (bottom) from the NAcSh. **f**, Summary data for saline-treated female D1-MSN vs D2-MSN neuronal excitability recorded during diestrus. **g**, Summary data for saline-treated vs cocaine-treated female D1-MSN neuronal excitability recorded during diestrus. **h**, Summary data for saline-treated vs cocaine-treated female D2-MSN neuronal excitability recorded during diestrus. **i**, Summary data for cocaine-treated female D1-MSN vs D2-MSN neuronal excitability recorded during diestrus. **j**, Representative raw traces from saline-treated and cocaine-treated female D1-MSNs (top), and D2-MSNs (bottom) from the NAcSh recorded during diestrus. **k**, Summary data for saline-treated female D1-MSN vs D2-MSN neuronal excitability recorded in estrus. **l**, Summary data for saline-treated vs cocaine-treated female D1-MSN neuronal excitability recorded in estrus. **m**, Summary data for saline-treated vs cocaine-treated female D2-MSN neuronal excitability recorded in estrus. **n**, Summary data for cocaine-treated female D1-MSN vs D2-MSN neuronal excitability recorded in estrus. **o**, Representative raw traces from saline-treated and cocaine-treated female D1-MSNs (top), and D2-MSNs (bottom) from the NAcSh recorded during estrus.

For females in diestrus, neuronal excitability differences under baseline conditions were similar (if not greater) to the pattern seen in male animals, i.e. reduced D1-MSN excitability when compared to D2-MSNs (Fig 2f). Interestingly, during estrus, excitability of D1-MSNs increased, to the extent that there were no differences between D1-MSNs and D2-MSNs (Fig 2k). Notably, the magnitude of change in D1-MSN excitability across the four-to five-day estrous cycle is comparable to the effect of cocaine in males (Fig 2k, 2b).

Cocaine administration to females produced additional unanticipated results. First, cocaine did not alter D1-MSN (Fig 2g) or D2-MSN (Fig 2h) excitability in females when measured during diestrus (Fig 2j). Second, during estrus, cocaine arrested D1-MSN plasticity (Fig 2l) while simultaneously initiating D2-MSN plasticity (Fig 2m). Specifically, there was an emergent increase in D2-MSN excitability (Fig 2o). In total, cocaine flipped the effect of the estrous cycle on D1-MSN/D2-MSN excitability. Under drug-naïve conditions, the estrous cycle balances D1-MSN/D2-MSN activity through increased D1-MSN excitability (Fig 2k), whereas following cocaine administration, the estrous cycle potentiated the discrepancy between the two subtypes of MSNs by enhancing D2-MSN excitability (Fig 2n).

The passive membrane properties of D1-MSN and D2-MSN across groups are shown in Table 1. As previously reported^22^,^23^, resting membrane potentials for D2-MSNs tended to be more depolarized compared to D1-MSNs, although this only reached significance in male and estrous females treated with saline.

**Table 1.** Passive NAcSh MSN membrane properties.

| | Capacitance<br>(pF) | $V_m$ (mV) | $R_m$ (MΩ) |
| --- | --- | --- | --- |
| Male D1-MSN saline (n=17) | 86.4 ± 3.5 | **-78.7 ± 0.9\*** | 104.1 ± 5.2 |
| Male D2-MSN saline (n=17) | 80.6 ± 4.7 | -75.7 ± 1.1 | 116.7 ± 7.7 |
| P value | 0.325 | **0.044** | 0.185 |
| Male D1-MSN cocaine (n=20) | 91.2 ± 3.4 | -77.9 ± 0.9 | 96.7 ± 4.9 |
| Male D2-MSN cocaine (n=18) | 84.6 ± 4.2 | -75.9 ± 0.8 | 111.8 ± 6.6 |
| P value | 0.223 | 0.111 | 0.07 |
| Female estrus D1-MSN saline (n=16) | 86.1 ± 3.5 | **-76.8 ± 0.6\*** | 94.1 ± 4.5 |
| Female estrus D2-MSN saline (n=13) | 88.9 ± 4.2 | -73.7 ± 1.0 | 95.8 ± 7.2 |
| P value | 0.608 | **0.007** | 0.838 |
| Female estrus D1-MSN cocaine (n=14) | 78.2 ± 3.1 | -75.6 ± 1.5 | **106.9 ± 4.6\*** |
| Female estrus D2-MSN cocaine (n=15) | 74.6 ± 4.0 | -73.5 ± 1.0 | 127.7 ± 8.2 |
| P value | 0.487 | 0.252 | **0.043** |
| Female diestrus D1-MSN saline (n=13) | 80.6 ± 3.4 | -76.2 ± 1.5 | 89.2 ± 4.8 |
| Female diestrus D2-MSN saline (n=14) | 83.3 ± 3.1 | -74.6 ± 1.1 | 101.4 ± 7.6 |
| P value | 0.603 | 0.394 | 0.193 |
| Female diestrus D1-MSN cocaine (n=14) | 81.3 ± 4.0 | -76.5 ± 1.0 | 92.9 ± 3.9 |
| Female diestrus D2-MSN cocaine (n=14) | 72.5 ± 3.2 | -73.5 ± 1.6 | 97.4 ± 7.9 |
| P value | 0.101 | 0.124 | 0.615 |

Abbreviations: $V_m$ = Resting membrane potential, $R_m$ = membrane resistance.

Given previous reports that cocaine can modulate glutamatergic synaptic activity in the NAcSh^8, 11, 12^, we next tested the effect of cocaine on mEPSCs. In males, we found enhancement of glutamatergic synaptic activity in cocaine-treated mice (Fig 3a-c). This was driven through increased mEPSC amplitude in both D1-MSNs and D2-MSNs (Fig 3c). For females, there were no estrous cycle effects in either mEPSC frequency or amplitude in saline-treated animals. For females recorded in diestrus (Fig 3d-f), cocaine caused enhancement of glutamatergic synaptic activity, solely through an increase in D2-MSN mEPSC amplitude (Fig 3f), although there was a strong trend for an increase in D2-MSN frequency (Fig 3e) as well. For females recorded in estrus, cocaine also enhanced glutamatergic synaptic activity (Fig 3g-i). This was produced by an increase in mEPSC frequency (Fig 3h) and amplitude (Fig 3i), but again, only in D2-MSNs. Finally, following cocaine treatment, an estrous cycle-dependent change developed in D1-MSN mEPSC amplitude (Fig 3f/i: hatched bars), with greater amplitudes (*p<0.0082) during estrus.

**Fig 3.**
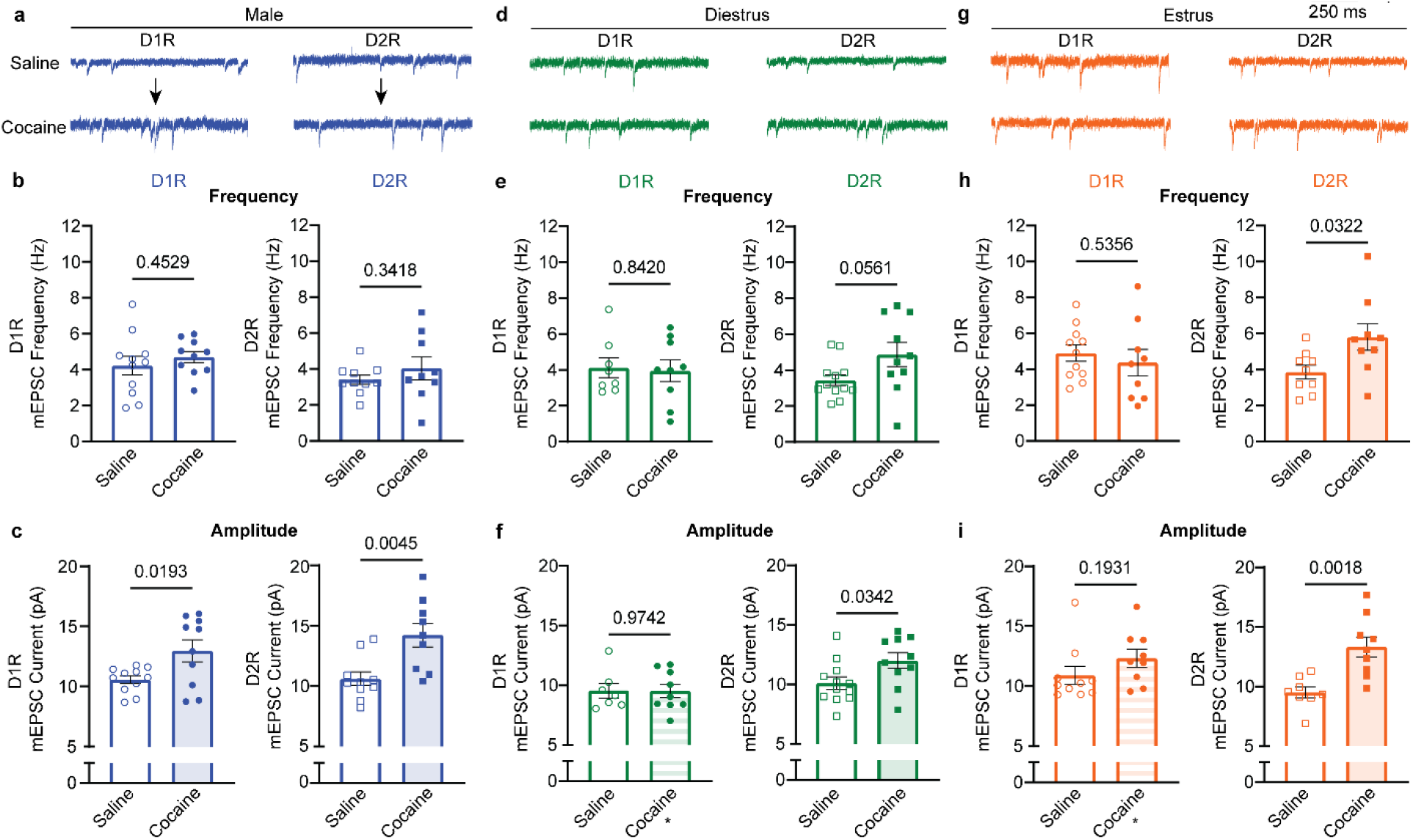
NAcSh mEPSCs in male and female mice. **a**, Representative mEPSC traces from NAcSh MSNs for saline and cocaine treated male animals. **b**, Summary NAcSh MSN D1-MSN and D2-MSN mEPSC frequencies for male animals treated with saline and cocaine. **c**, Summary NAcSh D1-MSN and D2-MSN mEPSC amplitudes for male animals treated with saline and cocaine. **d**, Representative mEPSC traces from NAcSh MSNs for saline and cocaine treated female animals recorded during diestrus. **e**, Summary NAcSh D1-MSN and D2-MSN mEPSC frequencies for female animals treated with saline and cocaine, recorded during diestrus. **f**, Summary NAcSh D1-MSN and D2-MSN mEPSC amplitudes for female animals treated with saline and cocaine, recorded during diestrus (*p<0.05 vs **i**). **g**, Representative mEPSC traces from NAcSh MSNs for saline and cocaine treated female animals recorded during estrus. **h**, Summary NAcSh D1-MSN and D2-MSN mEPSC frequencies for female animals treated with saline and cocaine, recorded during estrus. **i**, Summary NAcSh D1-MSN and D2-MSN mEPSC amplitudes for female animals treated with saline and cocaine, recorded during estrus.

This study is the first to explore the effects of cocaine on D1-MSN versus D2-MSN excitability of the NAcSh across sexes, with attention to the effects of estrous cycle prior to and after drug treatment. We have replicated previous findings that cocaine exposure leads to neuroadaptive effects in the NAcSh of male mice, primarily driven via an effect at D1-MSN^8–11, 13, 24–27^. This cocaine-induced D1-MSN driven plasticity is thought to be the primary contributor to the rewarding effects of the drug^28^ as well as the locomotor response^13^. Indeed in male mice, optogenetic stimulation of D1-MSNs of the nucleus accumbens mimicked cocaine reward, consistent with results in the dorsomedial striatum where D1-MSN optogenetic stimulation was deemed rewarding^29, 30^. Of notable contrast, D2-MSN stimulation in these same studies was found to be aversive.

Data from female animals implicate alternative hypotheses as to the role of D1-MSNs and D2-MSNs in cocaine action, or possibly alternative rationales for females versus male drug responsiveness. As some have hypothesized, it may be the gap or imbalance generated between D1-MSN versus D2-MSN activity in the nucleus accumbens^13^ which has a greater impact on the behavioral and rewarding effects of cocaine compared to strict D1-MSN modulation alone. As even with opposite adaptations (decreased D1-MSN excitability in males, increased D2-MSN excitability in females), the resulting disparity between D1-MSN and D2-MSN excitability following cocaine is quite similar. Alternatively, if D2-MSN activation is aversive in females as was observed in males, females might exhibit responses to cocaine more to minimize D2-MSN activation, a neuronal population that exhibits heightened excitability. This could be contrasted to males, which may be motivated to drive D1-MSN activation (which appears to be appetitive), from a population of neurons that exhibit decreased excitability.

A recent review hypothesized that males are more likely to engage in drug usage for sensation seeking (i.e. positive reinforcement), whereas females are more likely to take drugs as self-medication (i.e. negative reinforcement), downward spirals that are exacerbated with continued use^31^. It is provocative to consider that the sex differences reported here provide the neurophysiological underpinnings to support this hypothesis. Future experiments will further examine behavioral and physiological correlates regarding sex differences and drug-taking behavior to test this theory.

Another important finding is that D1-MSN plasticity is a normal process across the estrous cycle. Remarkably, the magnitude of change in D1-MSN excitability across the estrous cycle is comparable to the effect of cocaine in males (see Figure 2b,k). In females, cocaine arrested D1-MSN plasticity while simultaneously initiating estrous-cycle dependent changes in D2-MSN. The implications of these results are numerous. One, if D1-MSN and D2-MSN reflect appetitive and aversive responses as described above, motivational fluctuations across the estrous cycle are based on entirely different processes in drug na ïve versus drug treated individuals. Alterations in NAcSh physiology following cocaine administration are also enduring, at least in males^2,7^. It will be important to determine whether this shift in D1-MSN to D2-MSN plasticity in females across the estrous cycle endures not only for weeks, but possibly months or years. And while the mechanism by which cocaine produces these changes is currently unknown, it may also play a role in enhancement of cocaine conditioned place preference during estrus over diestrus^19^, the time where D2-MSNs exhibit heighted excitability.

Future studies will also need to determine whether other drugs of abuse produce similar sex-specific changes within the nervous system. As the rodent estrous cycle repeats every four to five days, emphasis should be placed in understanding how the hypothalamus-pituitary-gonadal axis produces these dynamic changes in neuronal activity. Provocatively, by having a greater understanding of the mechanisms by which the estrous cycle produces MSN plasticity, we may better identify treatments to reverse long-term drug-induced changes within these same populations of neurons across both sexes.

## MATERIALS AND METHODS

### Animals

Animal procedures were performed at the University of Minnesota in facilities accredited by the Association for Assessment and Accreditation of Laboratory Animal Care (AAALAC) and in accordance with protocols approved by the University of Minnesota Institutional Animal Care and Use Committee (IACUC), as well as the principles outlined in the National Institutes of Health *Guide for the Care and Use of Laboratory Animals*. Male and female mice were obtained from the Rothwell lab (University of Minnesota)^32^ and bred onsite. Mice aged 8-16 weeks, with a single copy of a Drd1a-tdTomato and/or Drd2-eGFP bacterial artificial chromosome (BAC) transgene were maintained on a C57BL/6J genetic background, group housed and kept on a 10:14 light:dark cycle with food and water *ad libitum*.

### Psychomotor sensitization

All experiments were conducted between 12:00 and 17:00, with houselights on at 06:00 and off at 20:00. Animals were handled and habituated to locomotor chambers as well as subcutaneous (S.C.) injections two days prior to behavioral testing. On test days, animals were given either an S.C. injection of cocaine (15 mg/kg) daily for five days ^2^, or an equivalent volume of 0.9% saline and placed promptly into the behavioral testing chamber (18” x 9”, with 8.5” tall walls) with light levels of 250 ± 10 lux. Videos were recorded for 80 minutes using ANY-maze tracking software (Wood Dale, IL) and total distance traveled analyzed offline.

### Chemicals

All chemicals were obtained from Sigma-Aldrich (St Louis, MO, USA), except cocaine hydrochloride which was obtained from Boynton Pharmacy (University of Minnesota, MN, USA).

### Whole-cell recordings

Mice (8-16 weeks old) were used for electrophysiology recordings. For females, prior to being anesthetized, estrous cycle was determined by vaginal cytology as previously described^33^. Mice were anesthetized with isoflurane (3% in O_2_) and decapitated. The brain was rapidly removed and chilled in ice cold cutting solution, containing (in mM): 228 sucrose, 2.5 KCl, 7 MgSO_4_, 1.0 NaH_2_PO_4_, 26 NaHCO_3_, 0.5 CaCl_2_, 11 d-glucose, pH 7.3-7.4, continuously gassed with 95:5 O_2_:CO_2_ to maintain pH and pO_2_. A brain block was cut including the NAcSh region and affixed to a vibrating microtome (Leica VT 1000S; Leica, Nussloch, Germany). Sagittal sections of 240 μm thickness were cut, and the slices transferred to a holding container of artificial cerebral spinal fluid (ACSF) maintained at 30 °C, continuously gassed with 95:5 O_2_:CO_2_, containing (in mM): 119 NaCl, 2.5 KCl, 1.3 MgSO_4_, 1.0 NaH_2_PO_4_, 26.2 NaHCO_3_, 2.5 CaCl_2_, 11 d-glucose, and 1.0 ascorbic acid (osmolality: 295–302 mosmol L^−1^; pH 7.3-7.4) ^4,34^ and allowed to recover for 1 hr. Following recovery, slices were transferred to a glass-bottomed recording chamber and viewed through an upright microscope (Olympus) equipped with DIC optics, a SOLA SE light engine (Beaverton, OR), the appropriate fluorescent filters, infrared (IR) filter, a fluorescence/IR-sensitive video camera (DAGE-MTI).

Slices transferred to the glass-bottomed recording chamber were continuously perfused with ACSF, gassed with 95:5 O_2_:CO_2_, maintained at room temperature and circulated at a flow of 2 mL min^-1^. Patch electrodes were pulled (Flaming/Brown P-97, Sutter Instrument, Novato, CA) from borosilicate glass capillaries with a tip resistance of 5–10 MΩ. Electrodes were filled with a solution containing (in mM) 135 K-gluconate, 10 HEPES, 0.1 EGTA, 1.0 MgCl_2_, 1.0 NaCl, 2.0 Na_2_ATP, and 0.5 Na_2_GTP (osmolality: 280–285 mosmol L^−1^; pH 7.3)^7^. D1-MSN and D2-MSN MSNs were identified under epifluorescence and, IR-DIC based on morphology and their hyperpolarizing membrane potential (−70 to −80 mV) and were voltage clamped at −80 mV using a Multiclamp 700B amplifier (Molecular Devices), currents filtered at 2 kHz and digitized at 10 kHz. Holding potentials were not corrected for the liquid junction potential. Once a GΩ seal was obtained, slight suction was applied to break into whole-cell configuration and the cell allowed to stabilize which was determined by monitoring capacitance, membrane resistance, access resistance and resting membrane potential (V_m_)^7, 35, 36^. Records were not corrected for a liquid junction potential of −15 mV. Cells that met the following criteria were included in the analysis: action potential amplitude ≥50 mV from threshold to peak, resting *V*_m_ negative to −66 mV, and <20% change in series resistance during the recording.

To measure NAcSh MSN neuronal excitability, V_m_ was adjusted to −80 mV by continuous negative current injection, a series of square-wave current injections was delivered in steps of +20 pA, each for a duration of 800 ms.

For miniature excitatory postsynaptic current recordings (mEPSC), slices transferred to the glass-bottomed recording chamber were continuously perfused with ACSF containing lidocaine (0.7 mM) to block voltage gated sodium channels and picrotoxin (100 μM) to block GABAR, and was continuously gassed with 95:5 O_2_:CO_2_, maintained at room temperature and circulated at a flow of 2 mL min^-1^. Patch electrodes were pulled (Flaming/Brown P-97, Sutter Instrument, Novato, CA) from borosilicate glass capillaries with a tip resistance of 5–10 MΩ and whole-cell recordings made.

Electrodes were filled with a cesium methanesulfonate (CsMeSO_4_) solution containing (in mM): 120 CsMeSO_4_, 15 CsCl, 10 TEA-Cl, 10 HEPES, 0.4 EGTA, 8.0 NaCl, 2.0 Na_2_ATP, and 0.3 Na_2_GTP (osmolality: 280–285 mosmol L^−1^; pH 7.3). D1-MSN and D2-MSN were identified under epifluorescence and, IR-DIC based on morphology and their hyperpolarizing membrane potential (−70 to −80 mV) and were voltage clamped at −80 mV using A Multiclamp 700B amplifier (Molecular Devices), currents filtered at 2 kHz and digitized at 10 kHz. Holding potentials were not corrected for the liquid junction potential. mEPSCs were recorded for 2 minutes and analyzed offline using Mini Analysis software (synaptosoft) with an amplitude threshold set at three times the noise level.

### Statistical analysis

Data values were reported as mean ± SEM. All statistical analyses were performed with a commercially available statistical package (GraphPad Prism, version 9.4.1). Probabilities less than 5% were deemed significant *a priori*. Depending on the experiments, group means were compared using a paired Student’s *t*-test, a one-way or a two-way ANOVA with repeated measures. Where differences were found, Bonferroni post hoc tests were used for multiple pair-wise comparisons.

## Acknowledgements

We would like to thank Dr. Erin Larson and Orion Rainwater of the UMN mouse behavior core for their help, Dr. Patrick Rothwell for providing the mice, proofreading and helpful suggestions, and Dr. Robert Miesel, Dr. Manuel Esguerra and Dr. Andréa Collins for proofreading and helpful suggestions.

## Author Contributions

A.D.C., C.A.N., P.P.J. and C.H.P. performed experiments; A.D.C. and C.A.N. analyzed data; A.D.C., C.A.N., M.J.T., P.G.M., prepared figures; A.D.C., C.A.N., M.J.T., and P.G.M., drafted manuscript; A.D.C., C.A.N., M.J.T., P.G.M. interpreted results of experiment; A.D.C., C.A.N., P.P.J. C.H.P., M.J.T., and P.G.M. edited and revised the manuscript.

